# Femtosecond pulsed laser ablation for in-tissue electron cryo-tomography

**DOI:** 10.64898/2026.10.28.761815

**Authors:** Jake L.R. Smith, Renae N. Ganon, Calina Glynn, James H. Naismith, Jorge Filevich, Michael Grange

## Abstract

*In situ* electron cryo-tomography (cryo-ET) is a powerful tool for visualising biology inside cells at the molecular scale. The routine application of cryo-ET in tissue samples promises biological insight beyond single cells, but is currently limited by the technical challenge of preparing high quality, electron-transparent specimens from thick (*>*100 µm) biopsies efficiently. Here, we demonstrate that cryogenic femtosecond pulsed laser ablation (cryo-fs-PLA) can accelerate the preparation of vitreous specimens of mammalian brain tissue for cryo-ET. Using cryogenic fluorescence microscopy, regions of interest within high-pressure frozen tissue can be specifically targeted, and cryo-fs-PLA can be employed as a preparative step for serial cryogenic lift-out experiments. We define cryo-fs-PLA parameters and determine ablation rates for two different laser wavelengths, and demonstrate that tilt series collected on lamellae fabricated from sites exposed to cryo-fs-PLA largely maintain their vitrification state. Incorporating cryo-fs-PLA into existing cryo-ET workflows has the potential to unlock the sample preparation of tissues at the throughput necessary for routine *in situ* cell biology of multicellular samples.

## Introduction

Mapping the molecular structure and organisation of protein assemblies directly within the cell using cryo-ET has provided vital insight into fundamental biological processes^1^. To visualise these processes *in situ*, thin (*<*200 nm) sections of cells amenable to transmission electron microscopy (TEM) must be first fabricated. For this, the implementation of focused ion beam (FIB) milling has been a crucial methodological development ^2,3^. With advancements in both hardware and automation, the generation of cellular lamellae for cryo-ET has become increasingly routine ^4–6^.

Beyond unicellular systems, cryo-ET can also be utilised in multicellular models to investigate the structural cell biology of tissues^7^. Typically, these samples cannot be vitrified by plunge-freezing on electron microscopy grids into a cryogen, therefore high-pressure freezing (HPF) in metal carriers is necessary to enable sufficient cooling rates for complete vitrification ^8,9^. Due to the increased sample dimensions and presence of the metal carrier, non-standard FIB-milling applications are required to make amenable samples for cryo-ET. In particular, the utilisation of cryogenic lift-out (cryo-lift-out) of HPF samples has been instrumental in enabling the structural cell biology of small organisms^10–14^, organoids^15^ and tissues^16–21^.

The excavation of large volumes of HPF material (i.e. trenches) by FIB-milling or cryogenic ultramicrotomy to isolate regions of interest (ROIs) from the bulk of the material is often a bottleneck for cryo-lift-out experiments. This step can be accelerated with the use of high milling current in both gallium FIB^12,16^ or inductively coupled plasma FIB (PFIB) milling ^17^. However, the process of material sputtering by FIB milling using high-current typically introduces surface artefacts and localised warming of ROIs near the milled area^22^. This requires subsequent “polishing” of the volume surfaces prior to lift-out using lower current FIB-milling, which significantly increases experimental time ^15^. Consequently, an approach that enables the rapid and routine removal of large volumes of material whilst preserving sample quality could transform structural cell biology of large (i.e. 0.1 mm to *>*3 mm) multicellular samples.

Pulsed laser ablation (PLA) offers a potential alternative to excavate large volumes of material efficiently. Using “ultrafast” pulses (i.e. on the femtosecond timescale), the removal of material proceeds faster than heat can be dissipated into the sample, and as such is considered an “athermal” process^23^. In addition, fs-PLA has been demonstrated to achieve 4–5 orders of magnitude faster than gallium and 2–3 orders of magnitude faster than PFIB milling in materials science samples^24^.

For beam-sensitive samples or liquid-containing samples, such as a Li-ion battery, fs-PLA can be performed at cryogenic temperatures (i.e. cryo-fs-PLA) which can permit the analysis of samples previously unattainable at room temperature ^25^. As maintaining the vitrification state of HPF samples is critical for cryo-ET analysis, the potential to reduce heat damage whilst decreasing processing time would be highly beneficial for in-tissue cryo-ET.

During the preparation of samples for analysis by other imaging or spectroscopic modalities, the increased throughput and reduced sample damage of fs-PLA relative to FIB-milling has been demonstrated in both materials science^26^ and life science^27,28^ applications. For example, fs-PLA has been utilised to prepare large pillars of chemically fixed, resin embedded mouse brain tissue for X-ray tomography imaging at scale^27^. However, the feasibility of integrating cryo-fs-PLA into the cryo-ET sample preparation workflow, where we benefit from making biological observations in a more native, hydrated state without the use of chemical fixatives, has yet to be performed. Additionally, the resulting effects of cryo-fs-PLA use on vitrified samples and their general quality, have yet to be described.

Here, we demonstrate how cryo-fs-PLA was able to prepare targeted cryo-lift-out sites from mouse olfactory bulb (OB) by the rapid removal of vitrified material with a minimal effect on the remaining sample. Our work therefore establishes cryo-fs-PLA as an effective preparative tool for tissue samples, and may thus both accelerate and simplify molecular level in-tissue structural biology.

## Results

### Integration of cryo-fs-PLA into the cryo-lift-out workflow for HPF tissue samples

A summary of the integrated cryo-fs-PLA/PFIB-milling workflow to target subfields in mouse OB with cryo-ET is presented in Figure 1. Using cryogenic fluorescence microscopy (cryo-FM), each subfield of the OB can be distinguished by the presence of neuronal cell body layers identified by nucleic acid staining prior to cryo-fixation (Figure 1a). Registration of the sample using SEM and FM overlays could be was performed to identify ROIs to prepare cryo-lift-out sites, and trench ablation using cryo-fs-PLA was performed adjacent to the ROI (Figure 1b). The resulting regions were subsequently isolated by cryo-lift-out (Figure 1c), where serial sections were deposited for lamella fabrication and subsequent tilt series acquisition (Figure 1d).

**Figure 1.**
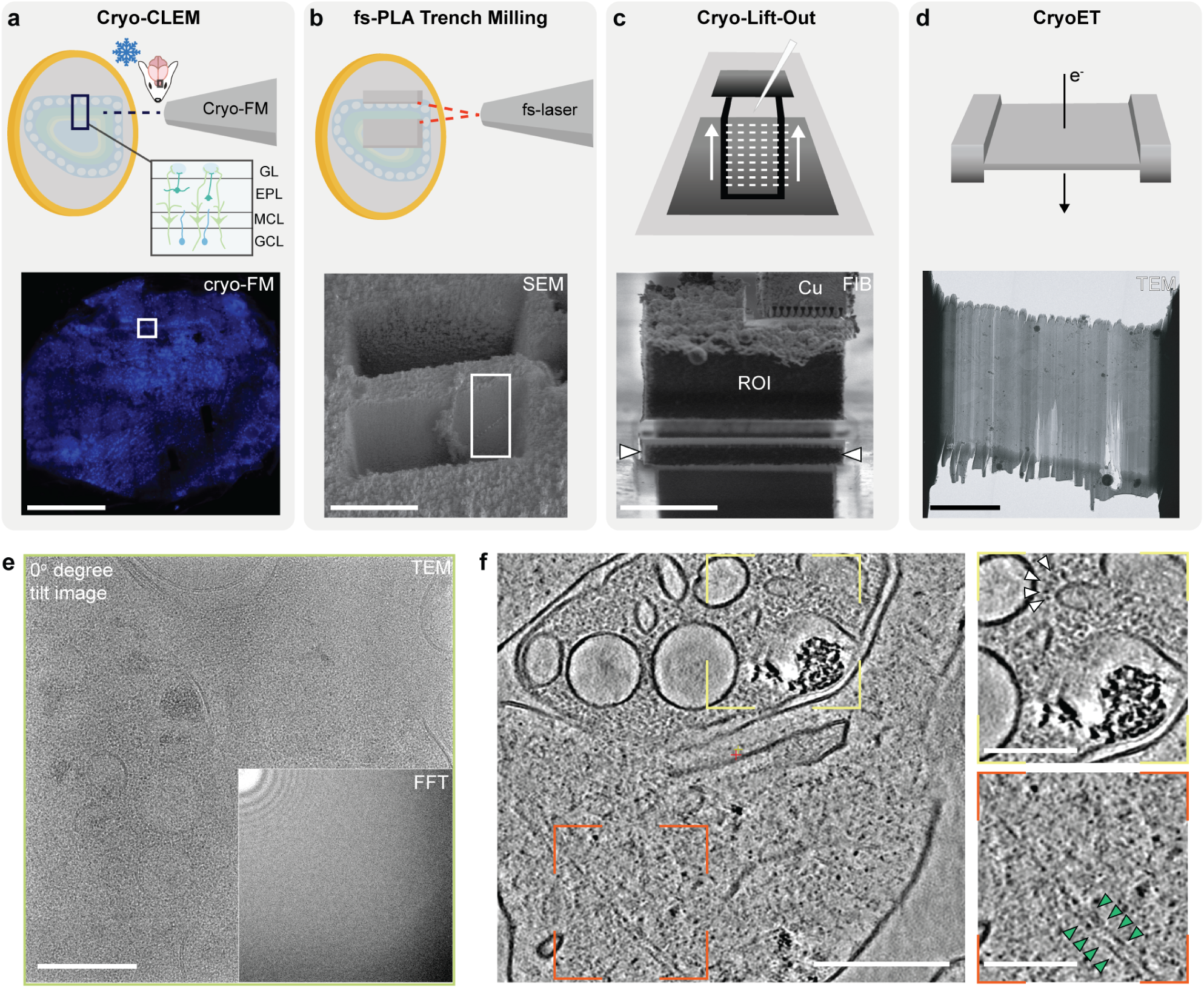
Workflow for fluorescence-targeted cryo-lift-out in mouse olfactory bulb. **a–d**, Depiction of the workflow for performing fluorescence-targeted cryo-lift-out in mouse olfactory bulb. Regions of high-pressure frozen tissue are targeted with cryo-fluorescence microscopy (**a**) before trenches are prepared with cryo-fs-PLA (**b**) and regions of interest (ROI) are identified (box). Serial cryo-lift-out is used to deposit sections of the ROI to a support grid (**c**) for lamella fabrication, followed by the collection of tilt series/cryo-ET data (**d**). **e**, TEM image of mouse olfactory bulb. The absence of crystalline ice reflections in Fourier space (bottom right) confirm that fs-PLA has no change on the vitrification state of the resulting lamellae. **f**, Representative Z slice of a resulting tomogram, showing mitochondrial ultrastructure (yellow) and filamentous actin (orange). Scale bars: **a**, 500 µm; **b**, 100 µm; **c**, 20 µm; **d**, 5 µm. **e–f**, 200 nm.

To prepare cryo-lift-out sites of HPF mouse OB using cryo-fs-PLA, the controlled ablation of material in a targeted manner is required. For this, a bespoke, dual-beam instrument equipped with a liquid nitrogen cooled stage was employed (Supplementary Fig. 1). For the ablation of trenches, the motorised stage could be tilted to orient the sample normal to the fs-laser beam. As the fs-laser was incorporated into a dual-beam instrument with an evacuated chamber, the sample can be positioned at coincidence with a scanning electron microscope (SEM) which allowed the ablation of material to be monitored *in situ* in real time.

### Ablation parameter definition for 515 nm and 343 nm wavelength fs-lasers

To assess the effect of performing cryo-fs-PLA on HPF samples, different ablation protocols were designed and applied to ablate trenches for cryo-lift-out. The speed of ablation and the surface quality of the resulting cross-section generated by cryo-fs-PLA can be controlled by changing the laser parameters (e.g. wavelength, pulse energy, pitch and repetition rate) ^29^. As fs-PLA is typically utilised for hard materials, protocols for the precise ablation of vitreous biological samples that produced trenches amenable to cryo-lift-out had to be established *de novo*.

Fs-lasers can be operated across the spectrum by utilising higher harmonics. Here, we tested a second harmonic green laser (515 nm) and a third harmonic UV laser (343 nm) for their performance on preparing HPF samples for cryo-lift-out. The focused spots for 343 nm wavelength pulses will be smaller than the spots for 515 nm wavelength pulses, meaning the energy density (fluence) of the 343 nm spots would be 2.25X larger than that for 515 nm spots (Supplementary Table 2).

For both 343 nm and 515 nm wavelengths, a “minimal” (lower pulse energy and larger pitch to target a “low dose” regime) and “optimal” (higher pulse energy and smaller pitch to target optimised cross-section surface and verticality) protocol were defined (Supplementary Table 3). Trenches of 150 × 150 × 100 were ablated with each of the wavelengths and protcols, taking between 16-131 seconds (Supplementary Table 3).

### Cryo-fs-PLA has a minimal effect on the vitrification state of lamellae

Prior to cryo-fs-PLA exposure, the vitrification state of two HPF OB biopsies was verified by TEM to ensure any observed non-vitreous artefacts were a result of sample-laser interaction rather than incomplete vitrification (Supplementary Fig. 2). The preparation of trenches with cryo-fs-PLA enabled a total of 14 cryo-lift-outs, leading to 29 lamellae and the collection of 412 tilt series. Of the 412 tilt series, 48 were removed prior to analysis due to bad tracking during collection or the lack of biological features (Supplementary Table 4).

To determine the effect of fs-PLA on the vitrification state, 364 tilt series were analysed for the presence of crystalline ice reflections in micrographs, and/or the appearance of reflections in Fourier space, indicative of the presence of cubic/hexagonal ice^8^ (Figure 2). In total, our data showed that 93.7 % of resultant tilt series displayed zero non-vitreous artefacts, with negligible difference in vitrification changes observed for both 343 nm (152/164, 92.7 %) and 515 nm (189/200, 94.5 %) protocols (Supplementary Table 4). The presence of non-vitreous artefacts may also be attributed to heterogeneous vitrification in certain regions of the sample during HPF, and not necessarily the effect of cryo-fs-PLA. These results suggest that the use of a fs-PLA approach does not alter the vitrification state of the sample in the vast majority of the collected data, and is thus compatible with *in situ* structural biology of tissues.

**Figure 2.**
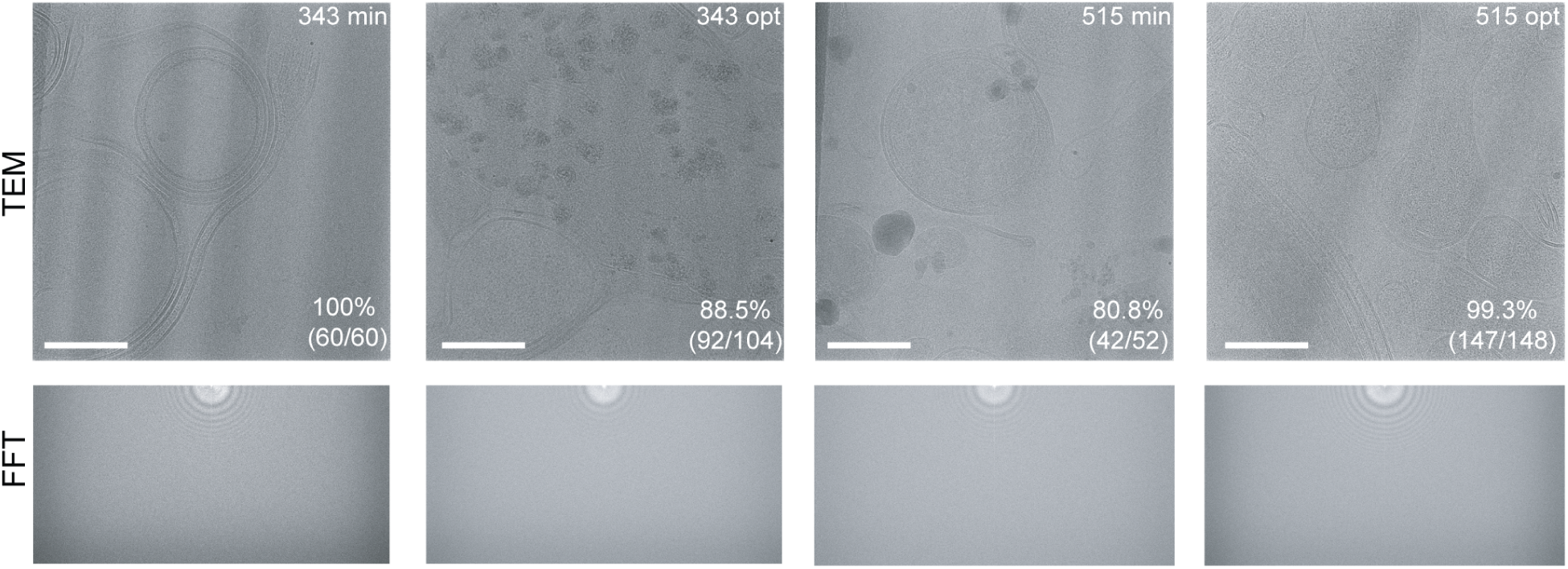
Assessment of vitrification state of cryo-fs-PLA prepared sections of HPF tissue samples. Representative cryo-TEM tilt images (**Top**) and the corresponding fast Fourier transform (FFT) (**Bottom**) from the four cryo-fs-PLA ablation protocols outlined in this study. Individual tilt series were analysed for the presence of non-vitreous artefacts to determine the percentage of vitreous tilt series. Scale bar: 200 nm.

One common observation in materials processed using fs-PLA is microstructural defects in the ablation-adjacent region ^30^. On closer inspection of TEM overview images, the edges of the lamellae adjacent to both 343 nm and 515 nm cryo-fs-PLA exposure showed the presence of sporadic hole artefacts (Supplementary Fig. 3a-b (Left)). Tilt series acquired in these regions did not show non-vitreous artefacts (Supplementary Fig. 3a-b (Right)).

To compare the artefacts between laser wavelengths, holes on lamellae obtained using 343 nm “min” (3 lamellae, 2 cryo-lift-outs) and 515 nm “opt” (5 lamellae, 2 cryo-lift-outs) were segmented for analysis. The area of the holes appeared slightly larger adjacent to 515 nm “opt” exposure (median 0.0035 µm^2^) in comparison to 343 nm “min” (median 0.0016 µm^2^) (Supplementary Fig. 3c). The median area per lamellae for both conditions overlapped, suggesting difference in artefact area may not be wavelength dependent (Supplementary Table 5).

Additionally, the hole artefacts appeared to propagate deeper into lamellae adjacent to 515 nm “opt” exposure (median 5.9 µm) when compared to 343 nm “min” (median 2.1 µm)(Supplementary Fig. 3d). All 5 lamellae generated adjacent to 515 nm “opt” had higher median distances from the exposure site (Supplementary Table 5). Due to the difference in ablation protocols for both wavelengths, and the small sample size (only a subset of lamella were suitable for analysis), the difference in artefacts cannot be attributed solely to the wavelength.

### Cryo-fs-PLA can remove large volumes of material rapidly in a targeted manner

To capture specific biological events within cells, correlative light and electron approaches (CLEM) can be performed which aid the preparation of cryo-lift-out sites within mouse brain tissue^17^. To demonstrate the compatability of rapid cryo-fs-PLA to target specific regions of HPF tissue for cryo-lift-out, large trenches of mouse OB were prepared which could be used to generate multiple adjacent cryo-lift-out sites (Figure 3a–b).

**Figure 3.**
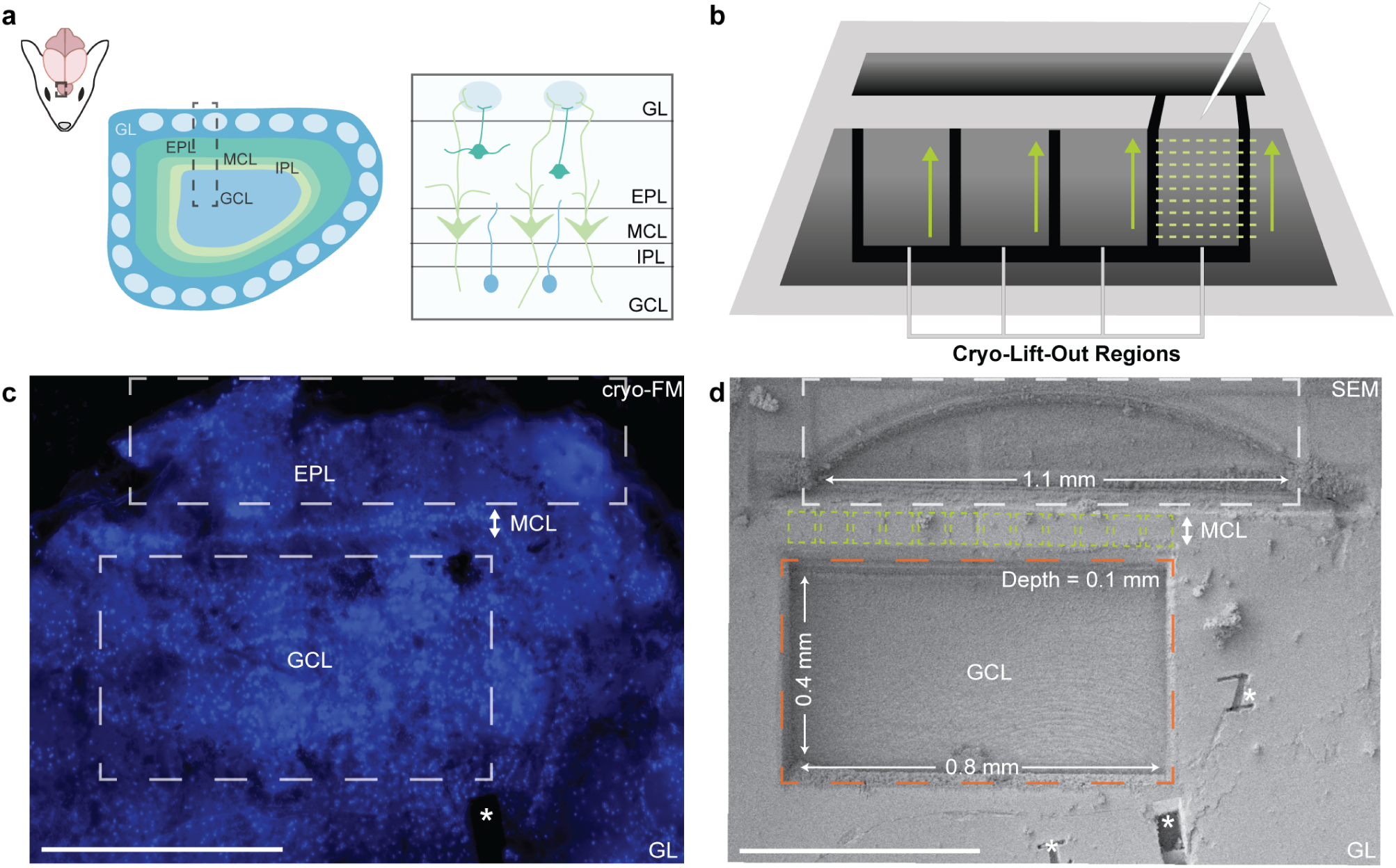
Demonstration of high-throughput cryo-fluorescence targeted trench milling in HPF mouse brain tissue. (**a**) Schematic overview depicting the workflow to obtain sections of mouse olfactory bulb. Right, diagram showing the different layers of mouse olfactory bulb (GCL, granule cell layer; IPL, internal plexiform layer; MCL, mitral cell layer; EPL, external plexiform layer; GL, glomerular layer). (**b**) Schematic depiction of the multiple adjacent cryo-lift-out sites (green) that can be prepared from large cryo-fs-PLA trench milling. (**c**) Cryo-FM of a high-pressure frozen mouse olfactory bulb section stained with Hoechst 33342 live-cell nuclear dye to identify subfields. Scale bar, 500 µm. (**d**) SEM image after cryo-fs-PLA of the region in **c** targeting the MCL, leaving a region of sample amenable to adjacent lift-outs (green boxes) as depicted in **b**. Scale bar: 500 µm.

As an example, for the fluorescence-guided cryo-lift-out of the mitral cell layer (Figure 3c), a single trench of 800 µm × 400 µm × 100 µm (total volume = 32 000 000 µm^3^) was excavated in just 144 s (ablation rate = 222 222 µm^3^ s^−1^) using cryo-fs-PLA operating at 343 nm (Figure 3d). In comparison to PFIB milling approaches, cryo-fs-PLA exhibits a ∼32-fold, ∼47-fold and ∼142-fold increase in ablation rate in comparison to 200 nA Xe^+^, 400 nA Ar^+^ and 120 nA Ar^+^ respectively (Supplementary Table 1). When compared to Ga^+^ FIB milling at 65 nA of vitreous ice, cryo-fs-PLA achieves a ∼444-fold increase^31^. This enhancement of site preparation efficiency could significantly reduce milling time for cryo-lift-out experiments, allowing previously inaccessible regions from deeper within tissue samples to be isolated rapidly for cryo-ET analysis.

## Discussion

Our results demonstrate that cryo-fs-PLA can be integrated into cryo-ET sample preparation workflows to accelerate the preparation of cryo-lift-out sites from HPF mouse brain tissue (Figure 1). The use of multiple wavelengths and protocols enabled the collection *>*350 tilt series with minimal effect on the vitrification state (93.7 % vitreous), showing the potential of cryo-fs-PLA for high-throughput cryo-ET studies of tissue samples. The ablation parameters established in this study, using two wavelengths (515 nm and 343 nm) each with two ablation protocols, provide the starting point in which further optimisation can be performed to improve cross-section quality, reduce artefacts and increase throughput ^29^.

As part of this work, we utilised a bespoke instrument integrating fs-laser optics and SEM into a “dual-beam” configuration (Supplementary Fig. 1). The incorporation of both modalities is analogous to existing commercial laser-FIB instruments ^24,26,32^, yet this is the first example of this configuration that permits the operation of fs-PLA at cryogenic temperatures with vacuum transfer loading and normal incidence laser processing. An advantage of this configuration compared with alternative approaches for isolating ROIs within thick HPF samples, such as the use of cryogenic ultramicrotomy to trim the metal HPF carrier ^15,33^, is that in principle, all steps from cryo-FM targeting (with integrated light microscopes in the FIB chamber^34^) through to cryo-lift-out could be contained within a single dual-beam microscope. This could minimise sample loss or damage via manual handling across multiple instruments.

With the use of cryo-fs-PLA, we demonstrate the ability to excavate large areas of sample at much higher rate than attainable with other FIB-milling approaches (Figure 3). For FIB-based cryo-lift-out, the trench milling step is typically where the majority of the instrument time is used due to the limits on the utility of high FIB milling currents. In comparison to existing FIB milling protocols that use high-current (i.e. over 65 nA), cryo-fs-PLA exhibits a substantial increase in trench ablation time (∼ 32-fold – 444-fold). By demonstrating that the preparation of sites can be performed much faster using cryo-fs-PLA (Figure 3), the ability to prepare both more complex samples (i.e. tissues) can be accelerated.

Finding ROIs within tissue samples requires a correlative approach, namely the use of cryo-FM to target labelled structures within cells^17^. Here, we demonstrate the cryo-fs-PLA can be used in conjunction with cryo-FM and cryo-SEM to ablate trenches in specific regions to target a large section of mitral cells within HPF mouse OB (Figure 3). As correlating ROIs in thicker HPF samples is more challenging due to the limited z-resolution of cryo-FM, additional steps to remove material to access features deeper within the sample is necessary. With cryo-fs-PLA, the increased ablation rate alongside the minimal change in vitrification state of the resulting sample could permit access to these regions, allowing more precise targeting for in tissue structural biology investigations.

Despite the vitrification state of the sample being largely maintained after cryo-fs-PLA, we observed the presence of holes at the lamella edge adjacent to the exposure site, which extend up to several microns into the sample (Supplementary Fig. 3). The extent to which these artefacts persisted into the sample appeared reduced for shorter wavelength lasers (343 nm “min”) than the longer wavelength pulse (515 nm “opt”) (Supplementary Fig. 3c). The sporadic nature of the artefacts could be due to the heterogeneous composition of the tissue sample, where the ablation rates of subcellular components (i.e. lipids, organelles) could cause material to be removed inconsistently. The mechanism by which these artefacts are formed is currently unknown; further work is required to understand their origin and to minimise them by optimising parameters such as pulse energy, wavelength, repetition rate, pulse overlap, etc. ^29^.

As the frontier in structural biology moves to in-tissue cryo-ET^7^, the application of cryo-fs-PLA in a correlative manner may prove critical in preparing ROIs within the deep regions of vitrified organoids and biopsies that are suitable in quality and number for effective cryo-ET investigations. Integrating cryo-fs-PLA into the cryo-lift-out workflow should expand the range of biological questions accessible by making lamella production from multicellular samples faster and more routine.

## Methods

### Animal Husbandry

All animal experiments were monitored and facilitated by Named Animal Care and Welfare Officers from the Mary Lyon Centre for Mouse Genetics at MRC Harwell, and animals were treated in accordance with the UK Animal Scientific Procedures Act (1986). The mouse used in this experiment was provided by the Mary Lyon Centre, Harwell, UK. A female adult (144 days old) C7BL/6J mouse (Jackson Labs, Bar Harbor, ME) was euthanised in a schedule 1 procedure via overdose with an intraperitoneal injection of dilute pentobarbital in buffered saline solution (1:1). Death was confirmed through permanent cessation of circulation via cutting of the femoral artery.

### Tissue Preparation for High-Pressure Freezing

The brain was dissected from the skull, and the olfactory bulb was isolated from the rest of the brain along with a small amount of cortex for manipulation. The tissue was then embedded in 3 % w/v low-melting point agarose (Thistle Scientific) in PBS by lowering the tissue into the agarose and cooling immediately on ice to set the orientation. The block was extracted and the excess agarose was trimmed before affixation to the vibratome chuck VT 1200S Vibratome (Leica Microsystems) and submerged in ice-cold dissection medium (99 % HBSS, 0.035 % w/v ascorbic acid, 0.002 % w/v ATP, 1 % v/v 100× penicillin/streptomycin solution, filter-sterilised through a 0.22 µm membrane, pH 7.1). 100 µm transverse slices were made by vibratomy set at frequency = 0.2 mm s^−1^, amplitude = 1.1 mm. The tissue sections were isolated from the surrounding agarose using curved tip needles (Bochem) and gently transferred to a well of a 24-well plate with resting media (300 µL Hibernate-A medium (Gibco) supplemented with 2 % v/v B-27 (Gibco)) for recovery and transport and kept on ice. Before freezing, slices were imaged by bright-field light microscopy (EVOS M7000, Thermo Fisher Scientific) to screen slice integrity and fluorescent staining quality.

### High-Pressure Freezing

High-pressure freezing carriers were prepared as described previously^35^. Briefly, the flat side of 3 mm type A gold-coated copper carriers (Leica Microsystems) were sanded with a combination of 4000/10000 grit sandpaper prior to metal polishing to ensure a smooth surface. Both the polished type A carriers and unpolished 3 mm type B 0.1/0.2 mm gold-coated copper carriers (Leica) were soaked in an excess of 1-hexadecene (Sigma-Aldrich) for at least 30 min before blotting dry with filter paper (Whatman).

After sectioning, slices were stained with Hoechst 33342 (Invitrogen) in the resting media. Slices were either taken whole or as 2 mm biopsies and submerged in cryoprotectant solution (NMDG-buffered artificial cerebrospinal fluid N-methyl-D-glucamine (NMDG) buffer (93 mM NMDG, 20 mM HEPES, 25 mM glucose, 2.5 mM KCl, 1.2 mM NaHCO_3_, 5 mM sodium ascorbate, 2 mM thiourea, 3 mM sodium pyruvate, 19 mM MgSO_4_(H_2_O)_7_, 0.5 mM CaCl_2_(H_2_O)_2_) with 10 % w/v sucrose (Sigma-Aldrich) and 10 % w/v dextran (MW 35–45 kDa) (Biosynth)) to incubate for between 26–33 min.

Slices were then placed inside the 0.1 mm recess of the pre-treated type B carriers, and excess cryoprotectant solution was wicked away using filter paper. The flat side of the pre-treated type A carriers were placed on top to close the freezing assembly before closing the lid and the initiation of freezing.

### Cryogenic Fluorescence Microscopy

Samples were screened using the bright-field fluorescence microscope (METEOR, Delmic) equipped with a 20× objective (NA 0.45) situated inside the chamber of a Helios 5 Hydra Dual Beam (Thermo Fisher Scientific) to visualise the layers of olfactory bulb in high-pressure frozen samples. Samples were placed into a 27°pre-tilted shuttle (Thermo Fisher Scientific) and loaded onto the stage of the dual beam. Fluorescence overviews were taken as described previously^17^. Briefly, multiple unique fiducial markers of ∼50 µm were milled into the surface of the sample (15 nA Xe, line pattern z = 0.5 µm) perpendicular to the beam for correlation between light and electron images. The centre of the sample was then positioned perpendicular to the fluorescence microscope where a tiled overview was obtained using Odemis software (Delmic). Images were acquired at 155 mW with 100 ms exposure, producing a total field of view of 1.88 mm × 1.87 mm.

### Cryogenic fs-Laser Configuration

Laser ablation experiments were performed using a prototype instrument (developed at Thermo Fisher Scientific, Hillsboro, OR, US). The configuration was based on a dual beam setup, integrated with a Verios 5 SEM column at coincidence with the fs-laser. A Carbide CB3-20W (Light Conversion) laser with 2nd and 3rd harmonic modules is situated in an interlocked, Class I-safe enclosure which is positioned externally to the microscope, and light is directed by mirrors via the injection port into the high-vacuum chamber in a similar way to the instrument described by Randolph et al. ^30^. Inside the chamber, shutters which protect the SEM during laser ablation are installed. The tool is equipped with a five-axis motorised stage, capable of 360*^◦^* rotation which enables alignment of ROIs. The stage is cryogenically cooled where HPF carriers can be mounted flat onto a sample shuttle and loaded into the chamber under vacuum. This allowed for the sample to be maintained at cryogenic temperatures at all times. The sample was positioned perpendicular to the fs-laser beam by tilting the stage for the ablation of trenches, which was controlled by custom control software^30^.

### Cryo-fs-PLA Parameters

HPF carriers containing vitreous mouse OB were loaded onto the cryogenically cooled stage and imaged with SEM, where registration of SEM and cryo-FM was performed using Maps 3.21 (Thermo Fisher Scientific) to guide targeting. Parameters for both 343 nm and 515 nm protocols were established iteratively by adjusting laser parameters, ablating regions of the sample and imaging the resulting cross-sections with SEM. Final parameters for each protocol were determined empirically based on the surface quality and the amenability of the trenches to be used for cryo-lift-out. These parameters are outlined in Supplementary Table 3.

### Cryo-Lift-Out Site Preparation

All cryo-lift-out experiments were performed on a Helios 5 Hydra CX PFIB-SEM (Thermo Fisher Scientific) equipped with a tungsten nanomanipulator needle with a copper block attachment. Trench milling for the pre-cryo-fs-PLA exposure was performed using PFIB milling. Additional trench milling for ablation rate comparison was performed using Ar^+^ (120 nA & 400 nA) and Xe^+^ (200 nA) without subsequent lift-out steps.

To isolate an ROI of 45 µm × 30 µm for lift-out, a front trench (60 µm × 100 µm) and rear trench (60 µm × 15 µm) were milled (60 nA, regular cross section (RCS)) by positioning the sample surface perpendicular to the PFIB (Stage Tilt = 27°, Stage Rotation = −110°). For the cryo-fs-PLA experiments, front and rear trenches (150 µm × 150 µm) were ablated using the protocols defined in Supplementary Table 3.

In a small subset (3/14) of cryo-lift-outs, rear trenches were produced using Xe^+^ milling (60 nA, RCS) to determine the directional effect of cryo-fs-PLA exposure from one side of the lamella. The large trench excavation depicted in Figure 3c–d was produced with a front trench (1900 µm × 300 µm front and 800 µm × 400 µm rear) using 343 “opt” with an 8 × 8 X-Y pitch (Supplementary Table 3).

### Cryo-PFIB Lift-Out

HPF samples and support grids were loaded under liquid nitrogen into a 27°pre-tilted life science shuttle (Thermo Fisher Scientific) before being transferred onto a cryo-cooled (−185 °C) stage.

Cryo-lift-out was performed using Xe^+^ plasma as previously described ^17^.Following a 180°stage rotation, a region of 45 µm × 50 µm was isolated from the bulk of the material by performing undercuts around the side (3 µm × 30–40 µm) and base (45 µm × 8 µm) using rectangle pattern (4–15 nA), leaving two small tabs attaching the region to the bulk of the material. Attachment and extraction of the region via the copper block attachment was performed by redeposition welding (0.3 µm × 1 µm × 5 µm, RCS, 1MP, 300 pA) followed by ablation of the connecting tabs (4 nA, rectangle) to release the sample for extraction.

### Serial Lift-Out

Serial sections were deposited on 400 × 100 mesh rectangular copper grids (Agar Scientific) pre-clipped into notched Autogrid rims (Thermo Fisher Scientific). The region was brought to the surface of the grid where a single weld (0.3 µm × 1 µm × 5 µm, RCS, 1MP, 300 pA) on both sides where the sample met the grid was performed. Sectioning was performed to create multiple sections 3–5 µm thick using a line pattern (1 nA Xe^+^) that spanned the diameter of the section. After depositing all sections, the needle was retracted and 3–5 additional welds (0.3 µm × 1 µm × 5 µm, RCS, 1MP, 300 pA) were applied to both sides of the sections using a ±20°stage rotation.

### Lamella Fabrication

Serial lift-out sections were thinned immediately after serial lift-out on the Helios 5 Hydra CX or loaded into the Arctis dual-beam PFIB-SEM microscope (Thermo Fisher Scientific) for lamella fabrication. Prior to milling, grids were sputter coated with platinum for 60 s at 12 kV and 70 nA, then coated with trimethyl(methylcyclopentadienyl)platinum(IV) for 70 s, followed by a further 60 s platinum sputter coating. Lamellae were thinned in an automated manner using AutoTEM Cryo 2.4 (Thermo Fisher Scientific), where sites were positioned on the top face of the sections to ensure the back of the section was retained in the final lamella. All milling was performed with Xe^+^ PFIB at 30 kV, with rough milling performed at 1 nA (2 µm offset, rectangle), 0.3 nA (0.7 µm offset, cleaning cross section (CCS), 1.5°over/undertilt), and 0.1 nA (0.3 µm offset, CCS, 1°over/undertilt). Polishing was performed at 30 pA (0.15 µm offset, CCS, 0.5° over/undertilt) and 10 pA (0.075 µm offset, CCS), and then manually polished until the protective platinum layer was lost. Final lamellae were sputter coated with an additional 10 s of platinum sputtering before transfer to the TEM to reduce charging.

### Cryo-ET Data Acquisition

Lamellae were loaded into a Titan Krios G4 (Thermo Fisher Scientific) operated at 300 kV. The microscope was equipped with: a stage capable of ±90°rotation; a Selectris energy filter (Thermo Fisher Scientific) operating with an energy slit width of 10 eV; and a Falcon 4i direct electron detector (Thermo Fisher Scientific). Tilt series were collected with off-axis beam shift with Tomo5 software (Thermo Fisher Scientific) using a dose-symmetric scheme from 60° to −60° from the lamellae fabrication angle in 3° increments. Tilt series were acquired as movies (.EER) at 64k× magnification (1.98 Å pixel size) at −5 µm defocus. The total dose per tilt series equalled 130 e^−^ Å^−2^.

### Tilt Series Processing and Tomogram Reconstruction

WarpTools^36^ was used for gain correction, motion correction and CTF estimation of tilt series data. Tomogram reconstruction was performed in AreTomo (version 1.2.5) ^37^ by weighted back projection with binning factor 4 (pixel size = 7.92 Å). Tomograms were postprocessed for visual inspection using CTF deconvolution and missing wedge correction in IsoNet2^38^. Tilt series and tomograms were visualised in 3dmod ^39^.

### Vitrification and Artefact Analysis

The vitrification of samples pre- and post-cryo-fs-PLA exposure was determined by visualisation in 3dmod ^39^. Each tilt image was assessed for non-vitreous artefacts in imaging and reciprocal space (via FFT), and a tilt series would be defined as “non-vitreous” if: ice reflections visible as dark patches, which persist through sample tilting were observed, or, the visible deformation of the cellular membranes. Tilt series were removed from the analysis if there was no observed biological contrast in the images, or if tracking between tilt angles was too poor to observe continuous features.

For the observed hole artefacts, segmentation was performed of TEM lamella overviews using the Segment-Anything-Models-Java (SAMJ-IJ) plugin for FIJI/ImageJ^40^, utilising the EfficientViT-SAM-I2 model ^41^ which reported the area of the holes. Coordinates for the centroid of each hole and the line at the lamella front were generated in FIJI^40^, and the distance between them calculated using a custom Python script. For each lamella, the median hole area and the distance are reported in Supplementary Table 5. Graphs were plotted in Python/Matplotlib ^42^.

## Acknowledgements

We would like to thank: Ms Maria Sanita and the Necropsy team at the Mary Lyon Centre at MRC Harwell for assistance with the preparation of animal tissue; Dr Matthew Case and Dr Tom Glen for operational assistance of the TEM and PFIB respectively; and Dr John Mitchels for facilitating this collaboration. This work was supported by the Electrifying Life Science project (220526/Z/20/Z to J.H.N.) and a Wellcome Career Development Award (225902/Z/22/Z to M.G.). J.L.R.S. was supported by a Wellcome Trust PhD Studentship (226810/Z/22/Z). The Rosalind Franklin Institute is funded by UK Research and Innovation through the Engineering and Physical Sciences Research Council (EPSRC). We acknowledge Diamond for access and support of the cryo-EM facilities at the UK national electron Bio-Imaging Centre (eBIC), proposal BI38637-8.

## Author Contributions

Conceptualisation – J.L.R.S., J.H.N., J.F. and M.G.; Investigation – J.L.R.S., R.N.G. and J.F.; Visualisation – J.L.R.S. and M.G.; Writing (original draft) – J.L.R.S. and M.G.; Writing (review & editing) – J.L.R.S., R.N.G., C.G., J.H.N., J.F. and M.G.; Supervision – C.G., J.H.N., J.F. and M.G.

## Competing Financial Interests

R.N.G. and J.F. were employed by Thermo Fisher Scientific at the time of this study. Thermo Fisher Scientific, The Rosalind Franklin Institute and the University of Oxford have an application pending on behalf of co-inventors J.L.R.S, R.N.G, J.F. and M.G. in the United States (US 19/281,921), Japan (JP 2026-176531), China (CN 202611126773.3) and Germany (DE102026131571.2) incorporating work described in this manuscript.

## Data Availability

The cryo-ET data generated in this study are available from the corresponding author upon request.

## Supplementary Information

**Supplementary Figure 1.**
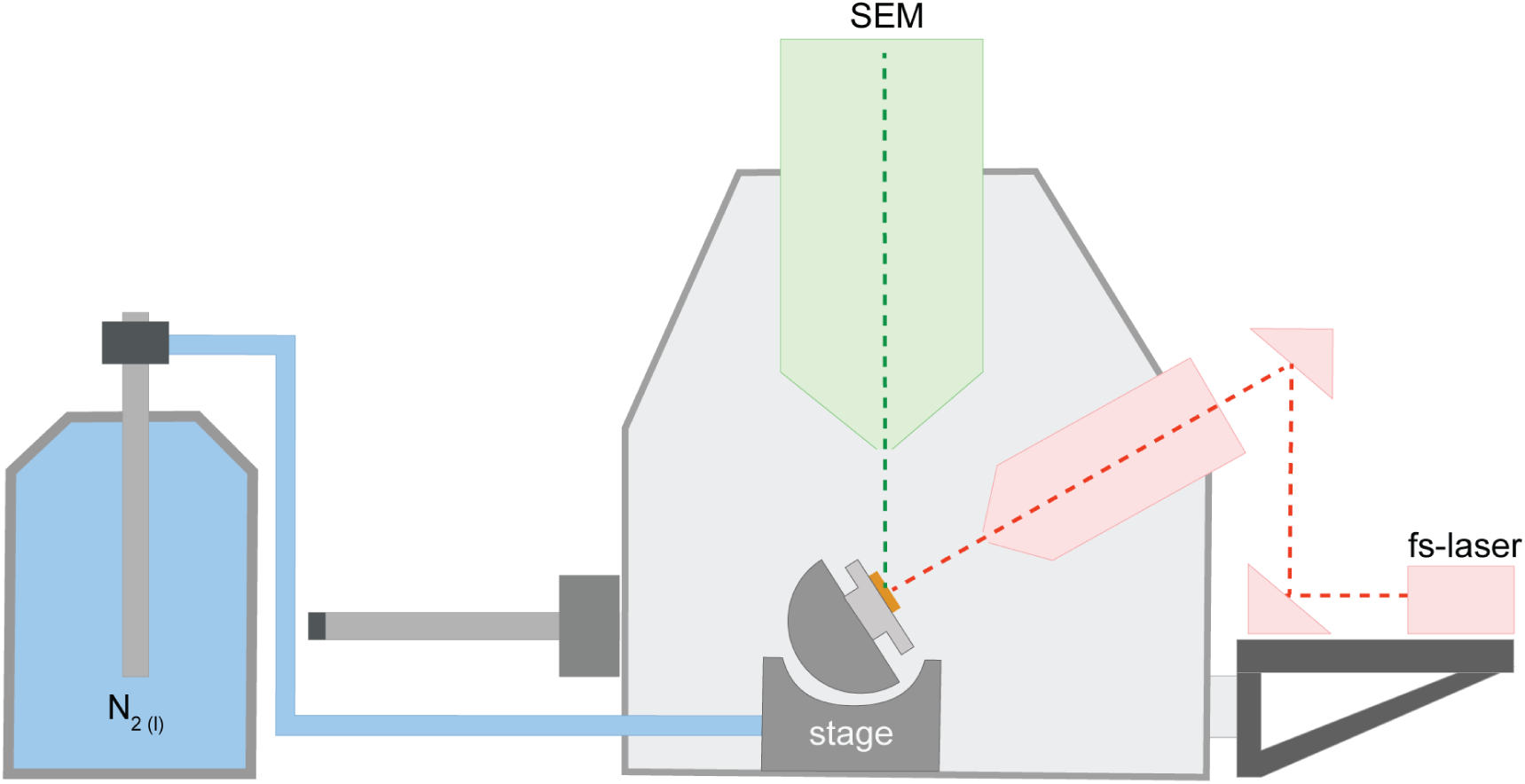
Schematic of the dual-beam configuration incorporating the fs-laser. A depiction of the dual-beam instrument that incorporates a scanning electron microscope (SEM) with a fs-laser. The two beams are at coincidence with the sample, which is loaded into the evacuated chamber via vacuum transfer, and held at liquid nitrogen temperatures

**Supplementary Figure 2.**
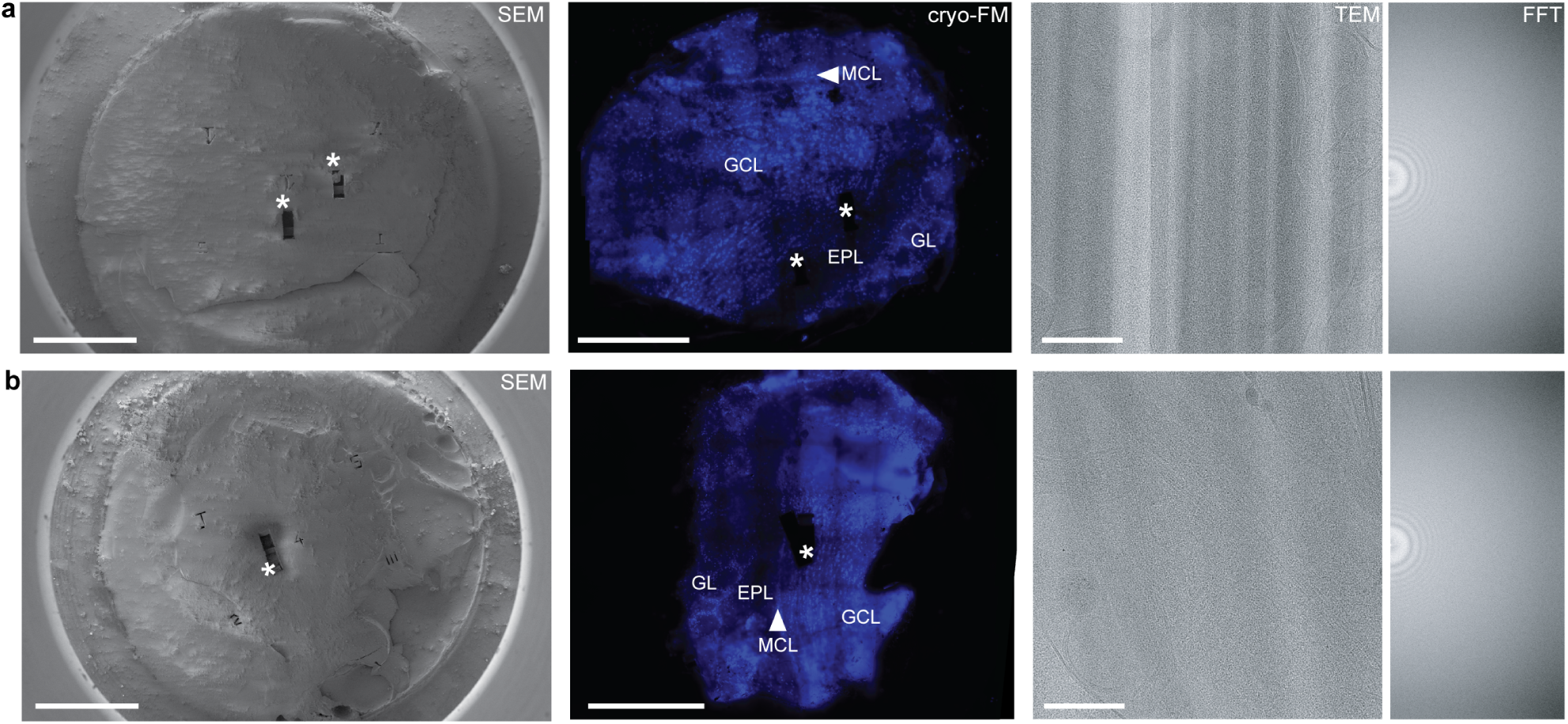
Vitrification screening for HPF tissue samples prior to cryo-fs-laser ablation. Two biopsies (**a-b**) of mouse olfactory bulb were high-pressure frozen, and their vitrification state was verified by cryo-ET. Corresponding SEM (**Left**) and cryo-FM (**Centre**) overviews were taken of the samples post-cryo-lift-out (*). Right, tilt series acquired from both samples show no non-vitreous artefacts. Scale bars: **Left - SEM**, 500 µm; **Centre - Cryo-FM**, 500 µm; **Right - TEM**, 200 nm.

**Supplementary Figure 3.**
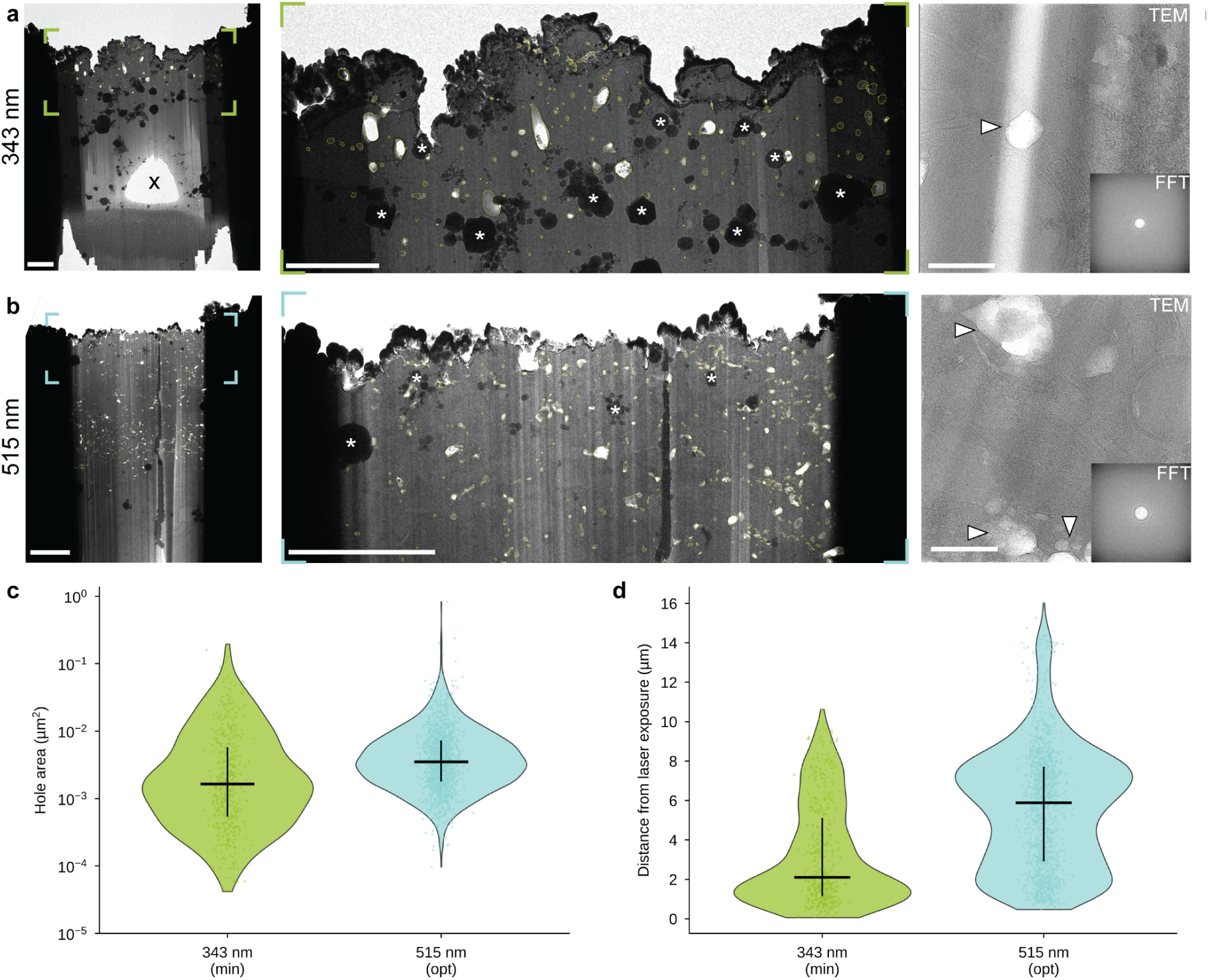
Assessment of effect of cryo-fs-PLA on the back edge of prepared lamella at different wavelengths. **a-b Left** TEM overviews of representative lamellae depicting the artefacts observed on the edges of lamellae adjacent to 343 nm“min” (**a**) and 515 nm“opt” (**b**) cryo-fs-PLA exposure. The cryo-fs-PLA was performed at the top of the lamellae, with the beam interacting with the sample perpendicular to the lamella. A tear from final lamella polishing is denoted by (x). A tear from final lamella polishing is denoted by (*). Scale bar, 1 µm. Centre, inset of **a** and **b** showing the segmented hole artefacts (yellow) at higher magnification. Dark black ice contamination from microscope transfer is denoted by (*). Scale bar, 1 µm. Right, tilt series collected in proximity to these artefacts show no non-vitreous artefacts. Hole artefacts present in the tilt series (black) do not change the vitrification state of the sample. **c** Comparison of the area of hole artefacts observed in lamellae generated from 343 nm“min” and 515 nm“opt” cryo-fs-PLA. **d** Comparison of the centroid distance of hole artefacts from the site of laser exposure for 343 nmmin and 515 nmopt. Both **c** & **d** are plotted with median (horizontal line) and interquartile range (vertical line) values.

## Supplementary Tables

**Supplementary Table 1.** PFIB milling parameters and total milling times for a 150 *×* 150 *×* 100 µm trench.

| Plasma | Pattern | Current (nA) | Total time (s) | Ablation rate ($\mu\text{m}^3 \text{s}^{-1}$) |
| --- | --- | --- | --- | --- |
| Xe | RCS | 200 | 322.5 | 6899 |
| Ar | RCS | 120 | 1422.5 | 1564 |
| Ar | RCS | 400 | 467.5 | 4759.3 |

**Supplementary Table 2.** Laser parameters and energy density correction for 343 nm and 515 nm “minimal” and “optimal” protocols.

| Wavelength (nm) | Protocol | Pulse energy ($\mu\text{J}$) | Spot size ratio | Area ratio | Correction factor | Corrected relative energy density |
| --- | --- | --- | --- | --- | --- | --- |
| 343 | Min | 3.5 | 0.666 | 0.444 | 2.25 | 7.89 |
| 343 | Opt | 4.5 | 0.666 | 0.444 | 2.25 | 10.14 |
| 515 | Min | 5.5 | 1 | 1 | 1 | 5.50 |
| 515 | Opt | 8 | 1 | 1 | 1 | 8.00 |

**Supplementary Table 3.** Parameters used for trench ablation using 343 nm and 515 nm “minimal” and “optimal” protocols.

| Wavelength (nm) | Protocol | Pulse energy ($\mu\text{J}$) | Repetition rate (kHz) | Max. average power (mW) | X/Y pitch ($\mu\text{m}$) | Ablation Time (s) for $150 \times 150 \times 100 (\mu\text{m})$ Trench |
| --- | --- | --- | --- | --- | --- | --- |
| 343 | Min | 3.5 | 1 | 3 | 5/5 | 43 |
| 343 | Opt | 4.5 | 1 | 5 | 3/3 | 131 |
| 515 | Min | 5.5 | 10 | 54 | 10/10 | 40 |
| 515 | Opt | 8 | 1 | 8 | 7.5/7.5 | 16 |

**Supplementary Table 4.** Assessment of vitrification state of cryo-ET data from cryo-fs-PLA trench milling protocols.

| Wavelength (nm) | Protocol | Tilt Series Collected | Tilt Series Analysed | Total Vitreous | Vitreous (%) |
| --- | --- | --- | --- | --- | --- |
| 343 | Min | 73 | 60 | 60 | 100.0 |
| 343 | Opt | 113 | 104 | 92 | 88.5 |
| 515 | Min | 55 | 52 | 42 | 80.8 |
| 515 | Opt | 171 | 148 | 147 | 99.3 |

**Supplementary Table 5.**
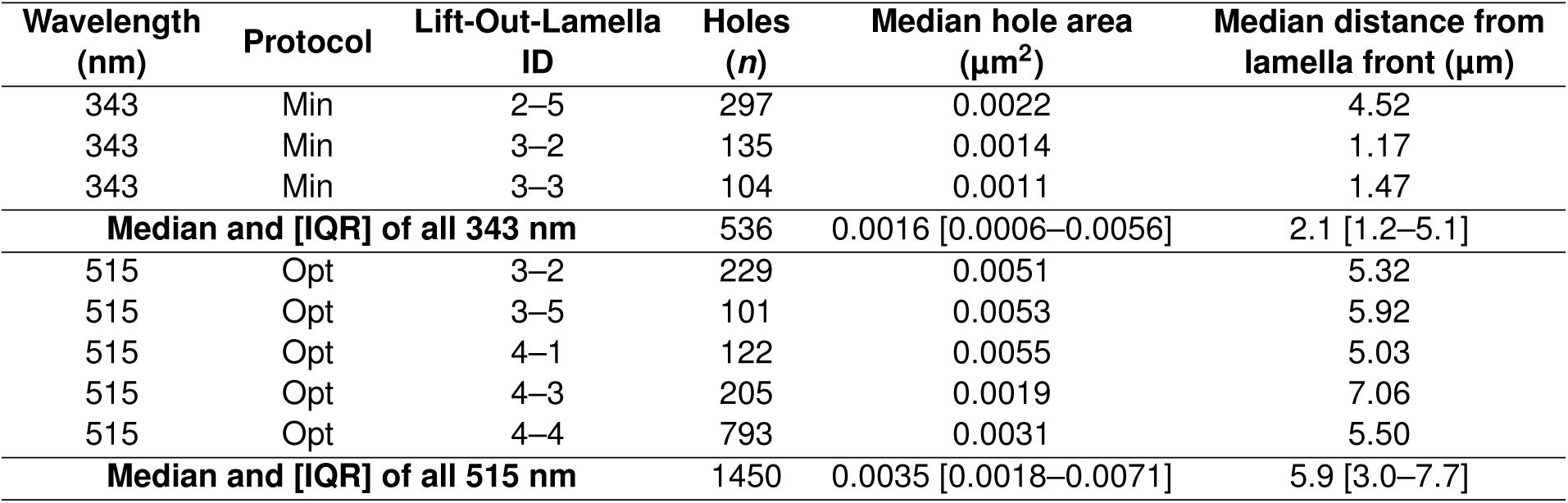
Hole artefact measurements for each lamella analysed using the 343 nm “minimal” and 515 nm “optimal” protocols.

| Wavelength (nm) | Protocol | Lift-Out-Lamella ID | Holes (*n*) | Median hole area (μm<sup>2</sup>) | Median distance from lamella front (μm) |
| --- | --- | --- | --- | --- | --- |
| 343 | Min | 2–5 | 297 | 0.0022 | 4.52 |
| 343 | Min | 3–2 | 135 | 0.0014 | 1.17 |
| 343 | Min | 3–3 | 104 | 0.0011 | 1.47 |
| **Median and [IQR] of all 343 nm** |  |  | 536 | 0.0016 [0.0006–0.0056] | 2.1 [1.2–5.1] |
| 515 | Opt | 3–2 | 229 | 0.0051 | 5.32 |
| 515 | Opt | 3–5 | 101 | 0.0053 | 5.92 |
| 515 | Opt | 4–1 | 122 | 0.0055 | 5.03 |
| 515 | Opt | 4–3 | 205 | 0.0019 | 7.06 |
| 515 | Opt | 4–4 | 793 | 0.0031 | 5.50 |
| **Median and [IQR] of all 515 nm** |  |  | 1450 | 0.0035 [0.0018–0.0071] | 5.9 [3.0–7.7] |

## Notes

### Summary of Updates

A revision to this manuscript was uploaded to correct the competing interests.

